# pmVAE: Learning Interpretable Single-Cell Representations with Pathway Modules

**DOI:** 10.1101/2021.01.28.428664

**Authors:** Gilles Gut, Stefan G. Stark, Gunnar Rätsch, Natalie R. Davidson

## Abstract

**Motivation:** Deep learning techniques have yielded tremendous progress in the field of computational biology over the last decade, however many of these techniques are opaque to the user. To provide interpretable results, methods have incorporated biological priors directly into the learning task; one such biological prior is pathway structure. While pathways represent most biological processes in the cell, the high level of correlation and hierarchical structure make it complicated to determine an appropriate computational representation.

**Results:** Here, we present *pathway module Variational Autoencoder* (pmVAE). Our method encodes pathway information by restricting the structure of our VAE to mirror gene-pathway memberships. Its architecture is composed of a set of subnetworks, which we refer to as pathway modules. The subnetworks learn interpretable latent representations by factorizing the latent space according to pathway gene sets. We directly address correlation between pathways by balancing a module-specific local loss and a global reconstruction loss. Furthermore, since many pathways are by nature hierarchical and therefore the product of multiple downstream signals, we model each pathway as a multidimensional vector. Due to their factorization over pathways, the representations allow for easy and interpretable analysis of multiple downstream effects, such as cell type and biological stimulus, within the contexts of each pathway. We compare pmVAE against two other state-of-the-art methods on two single-cell RNA-seq case-control data sets, demonstrating that our pathway representations are both more discriminative and consistent in detecting pathways targeted by a perturbation.

**Availability and implementation:** https://github.com/ratschlab/pmvae

## 1 Introduction

Transcriptomic analysis methods must find a balance between their ability to address the complexity of noisy molecular measurements, while simultaneously provide insight into complex biological processes. To gain biological insight into any high-dimensional molecular experiment, it is key to accurately identify and quantify the biological mechanisms underlying the observed changes across perturbations, cell states, or any other source of variability. A typical representation of underlying biological mechanisms are pathways; their structures remain mostly constant, but their changes in activity dictate a cell’s every action. Therefore, accurate estimation of pathway activity is a crucial aspect of the interpretation of single cell RNA-seq (scRNA-seq) data. Additionally, pathways have complex responses to perturbations, such as drugs, disease state, and environment. Identification of these effects on the expression level is challenging, since it is necessary to differentiate between effects due to perturbation versus inherent effects, such as cell type. This is further complicated by the fact that measurements may have multiple technical artifacts, such as batch effects and sparsity [16, 48]. Fortunately, deep learning techniques have successfully addressed many of the technical peculiarities of scRNA-seq data and may soon be integrated into typical bioinformatic workflows [32]. Their utility thus far has been demonstrated in tasks such as cell-type prediction [2], data harmonization [30, 35, 51], denoising [11], perturbation prediction [31] and visualization [35]. While effective, each of these models are black boxes and it is difficult to deduce from the model output what underlying biological mechanisms are driving the results.

A natural way to open up the black box of deep learning models is to interpret its learned parameters. For example, one can use variational autoencoders (VAEs) [21] on gene expression data and interrogate its latent representation. Way and Greene [52] correlate learned latent representations against external data, such as gene ontology terms and cancer subtype information, in order to explain the latent components. While this approach has proven fruitful [10, 23, 47, 53], it requires careful analysis to identify what each component is capturing, especially since all components are likely not fully disentangled [29] and thus could represent a combination of biological effects.

The use of the standard fully-connected layers in a deep learning model can introduce difficulties in interpretation. These layers lead to underspecified models in the sense that connections would exist between all pairs of genes, which allow the model to learn combinations of such biological effects based off of correlations not supported by our understanding of biology. One emerging approach has been to integrate prior information from biology, such as pathway gene sets, to constrain connections between entities known to interact [4, 13, 24, 33, 34, 38]. This integration of prior information not only aids in interpretability of the model but also its regularization.

Two approaches to integrate prior information in the form of pathways have been adopted: 1) represent each pathway as a gene set; or 2) represent the pathway as a hierarchical model. Models of the first approach take the form of factor analysis based methods that restrict latent factors to only explain genes within a pathway, as done in f-scLVM [4]. The Interpretable Autoencoder (interpretable AE) [38] is a modified hybrid between an auto encoder and a factor analysis model [45], by limiting the decoder to a linear projection and imbuing it with pathway information through regularization terms. While useful for finding overarching pathway changes, these approaches do not directly address the problem that many of these pathways are highly overlapping. PLIER [34] avoids this problem by clustering pathways through linear combinations of gene sets that are then represented as a single latent factor, but it is not intended to strictly model the activity of a single pathway.

The aforementioned overlap issue is significant when trying to quantify the downstream effects of a perturbed pathway. Genes that participate in a target pathway may also participate in downstream or adjacent pathways which introduce redundancies that are difficult to model. To take this effect into account, other methods have used hierarchical representations of pathways and their interactions. Similar to the flat representations, the methods first restrict the connections between the input layer and the initial hidden layer, in order to represent low-level pathway membership. After this first hidden layer, they connect hidden nodes to one another based upon a known pathway or biological process hierarchy. In DCell [33], GO annotations are used in order to hierarchically connect hidden nodes to represent the hierarchy of biological processes. In contrast, KPNN [13] uses the known signaling structure of signaling proteins, complexes, and families in order to represent the signaling cascades within a cell. To address additional difficulties in optimization due to redundancy within the hierarchy, DCell uses shortcut connections [26, 46], which also help optimize its large number of layers, and KPNN introduce a novel layer normalization scheme. While these work take pathway structure into account, they operate in a supervised learning domain, and learn representations suited for a prediction task.

An additional shortcoming of each pathway encoding scheme is that a single pathway is represented as unidimensional. Many higher-level pathways (e.g., the immune system) will contain possibly disparate signals from more specific and independent pathways (e.g., T-cell and B-cell signaling). This indicates that some pathways, like the immune system, require a richer representation than more specific pathways. The varying level of specificity naturally leads to the idea that some pathways can be represented by more than one dimension, and be reported in more detail than simply having increased or decreased activity. It has also been experimentally shown that some pathways are context dependent, which would require a multidimensional representation to fully capture its diversity [41, 54].

In this paper we present **pathway module VAE** (pmVAE), an unsupervised method to learn directly interpretable multidimensional pathway representations that take into account both pathway redundancy and context-specific activations. pmVAE incorporates pathways defined as a bag-of-genes, and constructs a latent space factorized by these pathways, resulting in multidimensional pathway representations. Through our pathway-module training procedure we are able to provide accurate pathway activities, even for highly correlated pathways.

## 2 Methods

pmVAE extends the VAE framework [21]. VAEs are probabilistic models that learn compressed representations of high dimensional data. They consist of two sets of functions, an encoder and a decoder, often parameterized by neural networks, with parameter sets *θ* and *φ* respectively. These models are optimized to learn distributions over low-dimensional latent variables, **z**, often referred to as embeddings or latent representations, from some high dimensional input data, **x**, by maximizing a lower bound on the log-likelihood of the data:

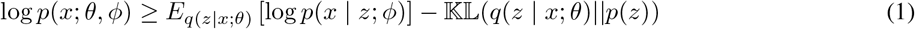

where 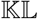 is the Kullback-Leibler divergence which regularizes the complexity of the embedding distribution. To make this optimization tractable, the posterior over latent representions, *q*(*z | x*) is often approximated with an isotropic Gaussian distribution and the prior over latent representations, *p*(*z*) is chosen to be a standard Gaussian. For a Gaussian likelihood *p*(*x | z*), the above optimization can be equivalently formulated as minimizing

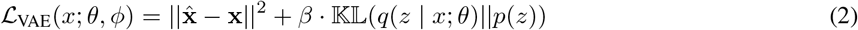

where the expectation is approximated at a single sample **z**, such that 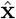 is the mean of the likelihood function evaluated at the drawn sample and implicitly depends on both *θ* and *φ*.

Optimization of VAEs are usually performed using online gradient based methods such as stochastic gradient descent, or popular second order methods, such as ADAM [20].

### 2.1 Pathway modules produce pathway specific representations

The pathway modules within pmVAE construct a latent space factorized by pathways. A graphic representation of the model is shown in Figure 1. Given a set of *K* pathways, each represented as a set of genes, pmVAE consists of *K* pathway modules, which each behave as a VAE constrained to the set of genes that participate in its pathway.

**Figure 1:**
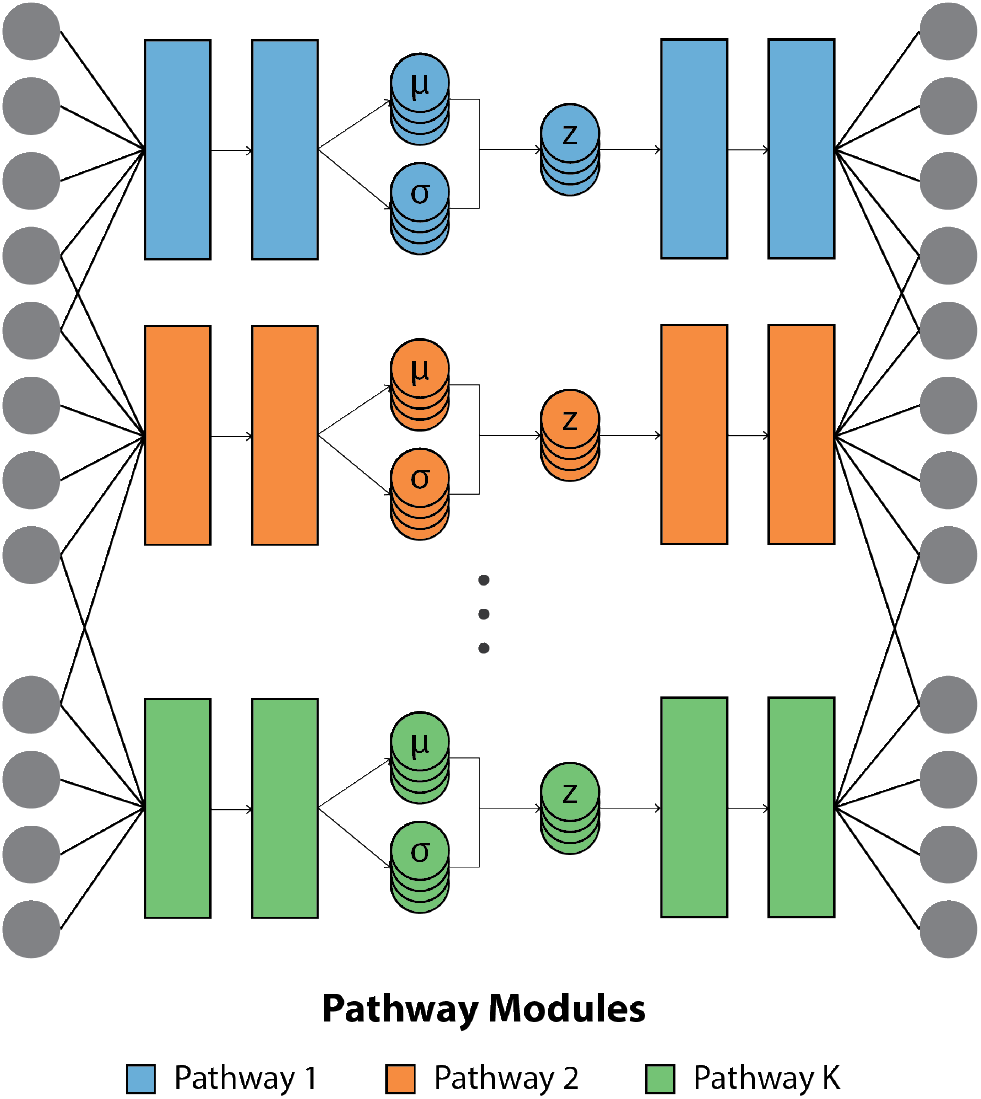
pmVAE is a variational autoencoder for expression data whose architecture incorporates prior knowledge about interacting genes using pathway modules. Pathways, defined as gene sets, are downloaded from community curated, public databases. Each pathway has a corresponding module which encodes and decodes only the genes participating within that pathway, producing latent representations and reconstructions specific to their pathway. Global reconstruction is achieved by summing over all pathway module outputs and a global latent representation of the input expression vector is achieved by concatenation of the latent representations from each pathway module. This constructs a latent space factorized by pathway where sections of the embedding explicitly capture the effects of genes participating in the pathway. We implement a custom training procedure to address optimization challenges caused by overlapping pathways, for example, due to hierarchical relationships. More details can be found in the methods section.

Let *N* be the number of total genes, *N*_*p*_ be the number of genes in pathway *p*, **x**^(*p*)^ be the expression of the genes participating in pathway *p* and let *θ*_*p*_ and *φ*_*p*_ be the parameters of the encoder and decoder within the pathway *p* module. Then the pathway *p* module encodes **x**^(*p*)^ into a pathway-specific embedding **z**^(*p*)^, which is then decoded into the reconstruction vector 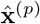. A global embedding vector, **z**is obtained by concatenation over all local embeddings provided by the pathway modules, i.e. *q*(*z | x*) = ∏_*p*_q(*z*^(*p*)^ | *x*^(*p*)^) and a global reconstruction is obtained by summing over the local reconstructions provided by each module by casting each 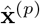 into a sparse vector with non-zero elements corresponding to the participating genes. This is achieved in practice by connecting the outputs of each module to the set of genes participating in the pathway (see Figure 1).

pmVAE minimizes the loss function:

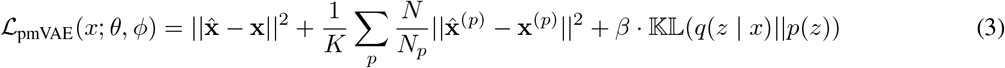

which consists of the usual global reconstruction and 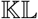 terms (the first and last of Equation 3), but also introduces a set of local reconstruction terms (middle). We additionally use an auxiliary pathway module that is connected to all genes in order to capture effects that could exist but are not contained within the ontology defining the pathway. This module is not included in the local reconstruction terms.

These local reconstruction terms enforce independence relationships between the modules, encouraging each module to independently reconstruct the genes participating in its pathway. Owing to the hierarchical nature of pathways, modules can exhibit some degree of redundancy, in the form of overlapping gene sets. This redundancy introduces degeneracies in the optimal solutions, causing problems in optimization. Due to the degeneracy, it can happen that one module is able to explain the effects of other overlapping modules, resulting in *XOR*-type behavior [13]. To see how this behavior arises, consider a naive loss function without the local reconstruction terms and a pathway gene set that is upstream from, and therefore overlapping with, a targeted pathway. If, through their mutual genes, the upstream pathway module is able to capture effects of the targeted pathway, then these effects will not be represented again in the targeted pathway, as this would result in a needlessly complex latent representation. The local reconstruction terms address this redundancy by enforcing independence relationships between modules.

We compute the local reconstruction terms in practice by performing an additional *K* gradient steps, each computed on the parameters of exactly one module, as shown in Figure 2. This process can be considered as some extreme form of the dropout regularization technique [42], in which all but one of the modules are dropped out in the forward pass. To prevent the model from favoring large pathways over small pathways, each local reconstruction term is weighted by size of the pathway relative to the global gene size 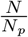. Furthermore, the 1 term introduces a trade off between an 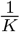 accurate global reconstruction and the independent local reconstructions.

**Figure 2:**
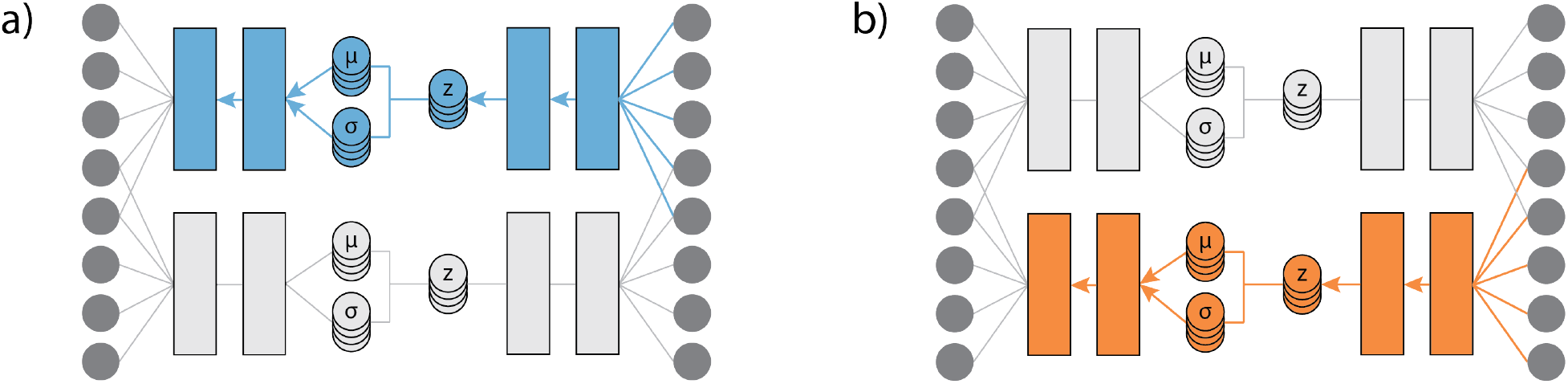
Visualization of the extra gradient steps in the training of pmVAE to compute the local reconstruction for the (a) first and (b) second module. In each step, gradients are not allowed to flow through the parameters of other modules, represented by greyed-out boxes. This procedure can be thought of as an extreme form of dropout [42] where all pathways except for one are dropped out. Due to the redundancies that exist between pathways, due in part to their hierarchical nature and cause optimization challenges, these extra gradient steps are necessary. If the redundancies are unaccounted for, it could happen, for example, that a single module explains the effects of a set of genes that mutually participate in other modules. By explaining these effects, this module effectively turns off the others, preventing them from capturing any effects [13]. This *XOR* behavior, i.e. where only a single module needs is needed to explain redundant effects seen across several modules, is a natural consequence of regularization, since a model that would re-capture these effects in multiple modules is needlessly complex. To alleviate this, these extra gradient steps are used to enforce independence between modules.

Finally, We make the usual Gaussian assumptions on the posterior and prior distributions over the embeddings and perform online optimization with the ADAM scheme [20].

### 2.2 Parallelization using dense masking layers

In order to take advantage of GPU-accelerated matrix multiplications, we define dense masking layers that parallelize the forward passes through all pathway modules. Theses masks are binary matrices which are element-wise multiplied with the usual kernel matrices of each layer in the network.

Two types of masks are used, defined below. Let *L* be the number of hidden layers, 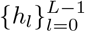 be the dimensions of the hidden layers within each module, *g* be the number of genes and *P*_*k*_ be the set of indices corresponding to genes participating in pathway *k*. The assignment mask *M* assigns genes to pathway modules and is defined as:

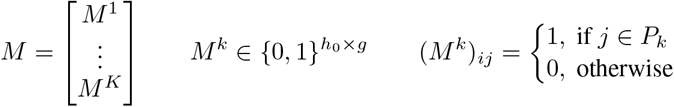

The separation masks 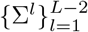 remove the connections between pathway modules at each hidden layer. They are block diagonal matrices defined as:

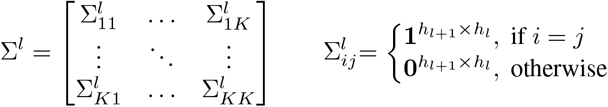

Let *ϕ* be the non-linear activation function and ⊙ be the element-wise matrix multiplication operator. For the *l*^*th*^ hidden layer let *W*_*l*_ and *b*_*l*_ be the usual kernel and bias terms, **y**_*l*_ be the output of the layer. Then the forward pass at each hidden layer (omitting the sampling step of the bottleneck layer) is defined as:

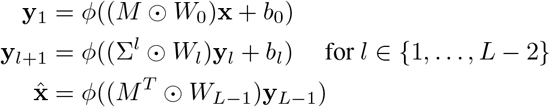

## 3 Data

To provide clear, quantitative comparisons against the other models, we wanted to identify datasets in which a perturbation with known downstream effects was applied. We consider two single cell RNA-seq case-control datasets from two independent sources where roughly half the cells were targeted with signaling molecules.

### 3.1 Pathway gene sets defined by Reactome [12]

We use pathway gene set annotations defined by Reactome [12] and maintained by MSigDB v4 (level C2) [44] in all experiments. The gene sets in Reactome are expert curated and hierarchically structured. There are a total of 674 gene sets with a median gene set size of 27 genes.

### 3.2 Interferon *β* stimulated cells from Kang et al. [19]

The Kang et al. dataset is a single cell dataset composed of peripheral blood mononuclear cells from eight lupus patients sequenced using droplet single-cell RNA-sequencing, with and without Interferon *β* stimulation. Preprocessing, in the form of library size normalization, removal of low variance genes and log scale transform, as well as pathway selection, was inherited from Rybakov et. al. [38], and yielded a dataset of 13,576 cells (7, 217 = 53.2% stimulated cells) by 979 genes, annotated with a total of 134 Reactome pathways, with a median gene set size of 21.

The signalling structure of pathways influenced by Interferon-*β* stimulation is summarized in Figure 3a. The interferon pathway is expected to be targeted as two of its child pathways are expected to change: Interferon-*α*/*β* signaling [36, 37] and `Antiviral mechanism by IFN-stimulated genes` [56]. We keep the suggestion of Rybakov et. al. [38] and consider four pathways to be the key pathways affected by the stimulation, which are among the parents and children of the targeted pathway in the Reactome hierarchy. As Interferon-*γ* and Interferon-*β* signal through two different receptors [36, 37], the Interferon *γ* pathway, which is one hierarchical level below, was not considered as an affected pathway.

**Figure 3:**
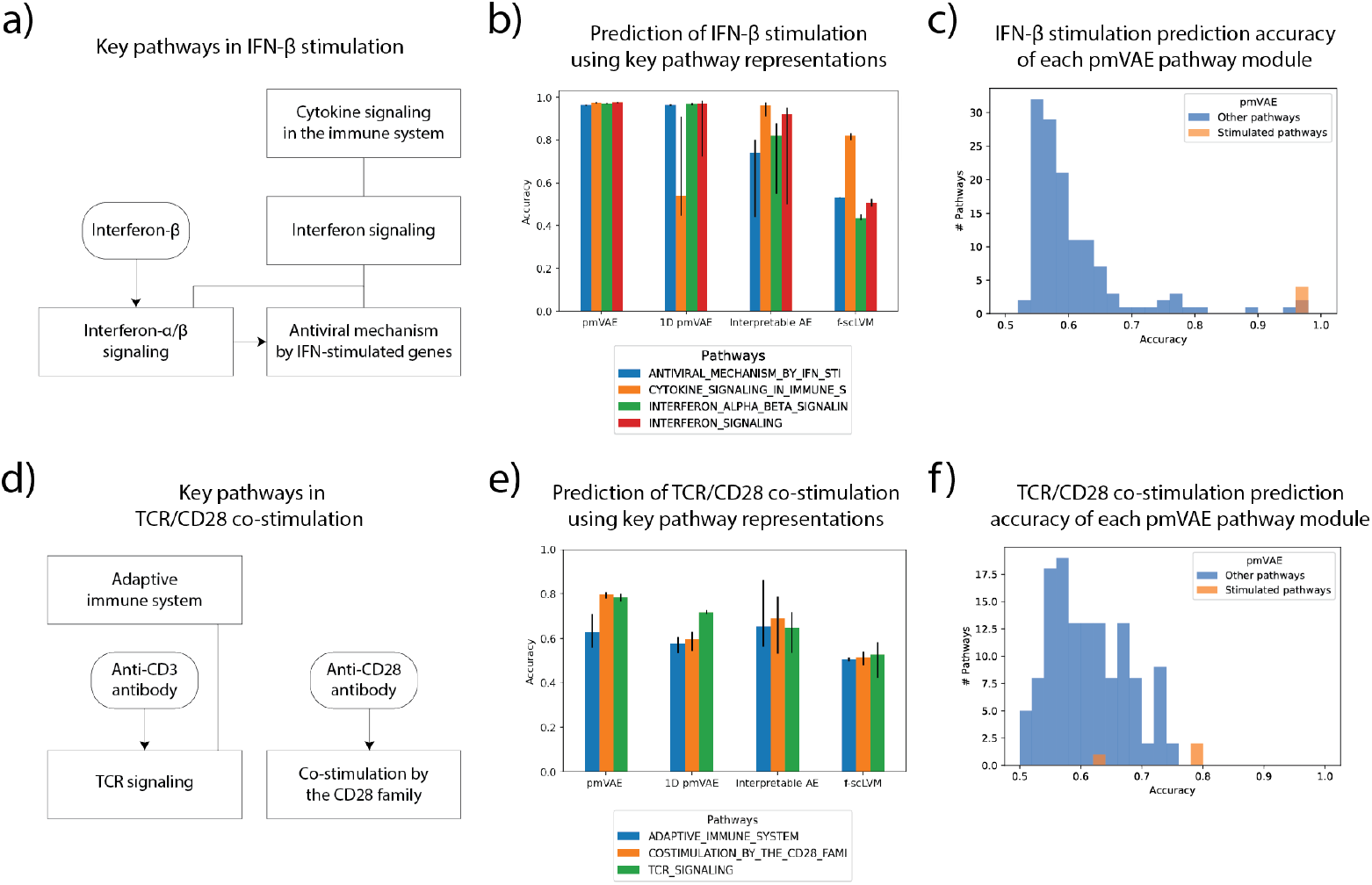
Stimulation targets and accuracy of pmVAE and baselines to predict stimulation status for the two considered experiments, top row: Kang et al. [19], bottom row: Datlinger et al. [7]. (a) and (d) Signaling structure, of the pathways targeted in the each experiment along their parents and children pathways, as defined by Reactome [12]. Square boxes represent pathways, curved boxes represent stimulation, lines between pathways represent hierarchical relationships and arrows represent activation events. For example, `Interferon-`*β* targets `Interferon-`*α*/*β* `signaling`, which targets `Antiviral mechanism by IFN-stimulated genes`, both of which are contained in `Interferon signaling`. (b) and (e) Logistic regression accuracies to classify perturbation status trained on pmVAE module embeddings and corresponding weights from baseline models, interpretable ae [38] and f-scLVM [4]. Error bars show the best and worst accuracy over 10 randomly seeded training procedures (10x cross-validation for f-scLVM). When considering the key pathways in each experiment, we observe that pmVAE consistently produces embeddings that are either as discriminative or better than baseline methods. (c) and (f) pmVAE logistic regression accuracies to classify perturbation status using the embeddings from each pathway module. The key pathways (orange) are among the most informative across all other Reactome pathways (blue). The most discriminative pmVAE modules are significantly enriched (P-Value < 0.05) in the top 5 pathways for the key pathways in each experiment.

### 3.3 TCR stimulated cells after guide RNA transduction from Datlinger et al. [7]

The Datlinger et al. dataset is a single cell dataset composed of Jurkat cells (immortalized human T-lymphocytes) that have been transduced with targeting and non-targeting gRNAs. Cells were starved or stimulated using anti-CD3 and anti-CD28 antibodies We only considered cells which had been transduced with non-targeting gRNA.

We filter genes that were expressed in less than 3 cells and filter cells with less than 200 expressed genes or greater than 10% mitochondrial RNA. We perform library size normalization using the 85^th^ percentile of each cell, rescale cells by a single randomly selected cell’s 85^th^ percentile, log-transform all values and rank genes by variance as described in Satija et al. [39] using the scanpy package [55]. Pathways with fewer than 15 genes in the top 5,000 highly variant genes are removed and genes participating in the remaining pathways are whitelisted from further removal. Finally, we remove low variance genes until we have 2,000 genes, that either participate in a retained pathway or pass the variance filter. This preprocessing procedure yields a dataset composed of 1,728 cells (853 = 49.6% stimulated) by 2,000 genes, annotated by 137 pathways with a median pathway size of 24 genes.

Since cells were targeted using anti-CD3 and anti-CD28 antibodies, we consider `TCR signaling` and `Co-stimulation by the CD28 family` to be the targeted pathways, the signaling structure is shown in Figure 3d. CD3 is a subunit of the TCR/CD3 complex [8] which transmits the activated TCR signal and enables the interaction of the T-cell receptor with other signaling molecules [3, 17, 27]. CD28 activation, together with TCR activation, results in co-stimulatory pathway activations [1]. We additionally considered all parent and child pathways, according to the Reactome hierarchy, of the two targeted pathways to be key pathways affected by the stimulation. From the set of retained pathways this includes `Adaptive immune system`.

## 4 Results

### 4.1 Hyper-parameters and model selection

For both datasets, pmVAE was trained using 1,200 epochs on 75% of the dataset. We initialize all kernels using uniform He initialization [14] and use ELU activations [6]. Multidimensional pmVAE modules use 4 latent dimensions per pathway. We perform a grid search over the learning rate (1e-2, 5e-2, 1e-3, …, 1e-5), *β* (1, 1e-1, … , 1e-8), and the architecture of the module encoders and decoders (one or two hidden layers with a width of 12 units). For the larger Kang et al. dataset we used a batch size of 256, and for the Datlinger et al. we used a batch size of 128. We perform model selection to choose parameters that lead to well-regularized models that did not need to sacrifice modeling power, by selecting the model with the largest *β* whose reconstruction error is within some tolerance (e.g., 15%) of the model with the minimum reconstruction error. For Kang et al. this selected a learning rate of 1e-3, *β* = 1e-5 and module encoders and decoders of a single layer and for Datlinger et al. this selected a learning rate of 5e-4, *β* = 1e-5, and module encoders and decoders of a single layer. On a single GeForce RTX 2080 Ti GPU, training pmVAE requires less than 4GB RAM and under 90 minutes for the Kang et al. dataset and under 30 minutes for the Datlinger et al. dataset.

We compare against two baseline methods, a linear factor analysis model, f-scLVM [4] and a hybrid factor analysis model, Interpretable Autoencoder (interpretable AE) [38], which is an auto encoder that has a linear decoder that reconstructs expression signals as a linear combination over gene sets defined by pathway membership. PLIER [34] was not considered because its latent variables cluster pathways instead of providing a single pathway score.

For interpretable AE, we inherit the parameters selected on the Kang et al. dataset. For the analysis of the Datlinger et al. dataset, we perform a grid search over the learning rate (1e-2, 5e-3, 1e-3, …, 5*e* 5) and the parameter *λ*_0_ (1, 1e-1, … , 1e-8). Using the reconstruction error for model selection, we select a learning rate of 0.005 and *λ*_0_ = 0.01. For f-scLVM, we use the suggested 3 dense factors and its deterministic optimization scheme.

### 4.2 Multidimensional pathway representations outperform unidimensional ones in identifying underlying biological effects

To demonstrate that increasing the dimensionality of a pathway leads to more accurate representations, we first compare 4-dimensional pmVAE, against the two unidimensional baseline methods, f-scLVM and interpretable AE, as well as a unidimensional pmVAE (1D pmVAE). We apply each method to two scRNA-seq datasets, Datlinger et al. [7] and Kang et al. [19]. Data preprocessing details are provided in Sections 3.2 and 3.3. In this task, we would like to demonstrate that the target and directly related pathways discriminate between the perturbed and control cells. Afterward, we learn a logistic regression model to predict stimulation status using the trained embeddings and compute the accuracy of this model on the yet unseen test data.

In Figure 3e we see that pmVAE is able to achieve high accuracy in the perturbed pathway, `TCR signaling` (accuracy: 0.78), and the other two related pathways (`Adaptive immune system` accuracy: 0.63, `Co-stimulation by CD28` accuracy: 0.80). Key pathway selection criteria are described in Section 3.1. When comparing the unidimensional methods, we find that the 1D pmVAE has significantly higher accuracy than the interpretable AE when using `TCR signaling` (1D pmVAE accuracy: 0.72; interpretable AE accuracy: 0.64; Mann Whitney U test p-value: 5.6 · 10^*−*4^) Interestingly, 1D pmVAE has decreased performance for pathways that are more distant to the perturbed pathway. In comparison, interpretable AE has the best performance when using the most distant upstream pathway, `adaptive immune system`. When considering only this pathway, we find that 1D pmVAE is outperformed, though not significantly (pmVAE accuracy: 0.63; interpretable AE accuracy: 0.66; Mann Whitney U test p-value: 0.1). The f-scLVM model has lowest accuracies across all pathways of interest. Top ten most accurate pathways for each model is given in the Supplemental Tables.

After demonstrating that pmVAE outperforms 1D pmVAE and interpretable AE in the target pathway on the Datlinger et al. dataset, we applied all methods to the Kang et al. dataset. Following the same experimental structure introduced in [38], we used Reactome as our pathway annotation with the key pathway being `Interferon` *α/β* `signaling`, with the following associated pathways: `Cytokine signaling in the immune system, Interferon signaling`, and `Anti-viral mechanism by interferon-stimulated genes`. Figure 3b reveals the performance for each method across the target pathway, `Interferon` *α/β* `signaling` and related pathways. We find that pmVAE matches or has greater performance in discriminating stimulated cells in comparison to all methods across all pathways. Again, the top ten most accurate pathways for each model are given in the Supplemental Tables. Both pmVAE and 1D pmVAE significantly outperform all other methods using the `Interferon` *α/β* `signaling` representation (pmVAE accuracy: 0.97; 1D pmVAE accuracy: 0.97; interpretable AE accuracy: 0.76; f-scLVM accuracy: 0.44). The only pathway in which 1D pmVAE is outperformed by the other models is `Cytokine signaling in the immune system`. This pathway is the most upstream pathway of all considered, meaning that this pathway must take into account the most downstream effects. Since the pathway activation in the 1D pmVAE can only be represented unidimensionally, we believe its representation is confounded by changes in other pathways. Overall, we observe that performance is greatly increased once the number of dimensions is increased.

### 4.3 pmVAE achieves the highest accuracy in detecting cell stimulation status using key pathway representations

To quantify the sensitivity and specificity of our pathway score, we applied pmVAE again on the Kang et al. and Datlinger et al. datasets. In both datasets, we aim to identify that the target and directly related pathways best discriminate between the perturbed and control cells in comparison to irrelevant pathways.

Figure 3c displays our performance on the Kang et al. dataset on the same task as described in Section 4.2 across all Reactome gene sets (134 in total) that have at least 12 genes. This figure demonstrates that pmVAE is well calibrated: all key pathways, shown in orange, are found to have the highest discriminative power when compared against irrelevant pathways, shown in blue, (median enrichment score using the top 5 pathways: 1.00 · 10^−4^; test: hypergeometric). Interpretable AE also achieves a significant enrichment score when considering only the top 5 most discriminative pathways (median p-value: 1.00 · 10^−4^; test: hypergeometric), but f-scLVM does not (p-value: 0.14; test: hypergeometric).

When we consider the Datlinger et al. dataset shown in Figure 3f, we again find that the most relevant pathways have the highest discriminative power (median enrichment score using the top 5 pathways: 3.15 · 10^−3^; test: hypergeometric). Neither interpretable AE nor f-scLVM have significant enrichment within the top 5 most discriminative pathways (interpretable AE median p-value: 0.10; f-scLVM p-value: 0.89; test: hypergeometric).

Furthermore, we sought to demonstrate that pmVAE does not suffer from module degeneracy, where strongly discriminative models can cause modules that contain overlapping genes to become less discriminative even though they are affected by the perturbation (see Section 2.1). Therefore, we would like to show that all relevant pathways have a discriminative ability comparable to the most discriminative pathway. We demonstrate this quantitatively by computing the average of the distance to the top rank among key pathways. The median value across all seeds is summarized in Table 2 and we find that pmVAE has the smallest values for both datasets.

**Table 1:** Median P-values from the hypergeometric test using the top 5 ranked pathways for the different methods and the two datasets over the 10 randomly seeded run. The hypergeometric test determines if the targeted pathways are overrepresented in the top 5 most accurate pathways in identifying perturbed cells. If the targeted pathways are also the accurate pathways, this shows that the model correctly identifies the relevant pathways to the perturbation. We see that pmVAE is the only method that is significantly enriched (P-value < 0.05) with targeted pathways in both datasets.

|  | Datlinger et al. | Kang et al. |
| --- | --- | --- |
| pmVAE | 3.15e-03 | 1.00e-04 |
| Interpretable AE | 0.10 | 1.00e-04 |
| f-scLVM | 0.89 | 0.14 |
| 1D pmVAE | 0.10 | 1.00e-04 |

**Table 2:** Median average rank difference to top ranked pathway. A high score here means that there is high variablity in the discriminative ability of the pathways, indicating that the modules could be degenerate. pmVAE has low scores for each experiment.

|  | Datlinger et al. | Kang et al. |
| --- | --- | --- |
| pmVAE | 17.5 | 2.0 |
| Interpretable AE | 19.5 | 16.4 |

### 4.4 Multidimensional pathway embeddings capture nuanced and context specific effects

In Sections 4.2 and 4.3, we quantitatively demonstrated that our multidimensional pathway representations are more discriminative of the underlying biological stimulus. In this section, we seek to demonstrate that these representations capture relevant biological signals that are important to a pathway. To more closely interrogate the pathway signals learned and show the benefit of the multidimensional representation, we use embeddings learned from the Kang et al. dataset to visualize various pathway representations. We do so by considering four pathways and their activation within different cell contexts. Two of the considered pathways are affected by the stimulation, `Interferon` *α/β* `signaling` and `Cytokine signalling in the immune system`. We also consider two pathways unaffected by the stimulation, `TCR signaling`, which is specific to T-cells, and a control pathway Cell cycle, which we expect to capture neither stimulation not cell type effects. We visualize the embeddings of these modules by computing tSNE projections [50] on each of the 4-dimensional embeddings, shown in Figure 4.

**Figure 4:**
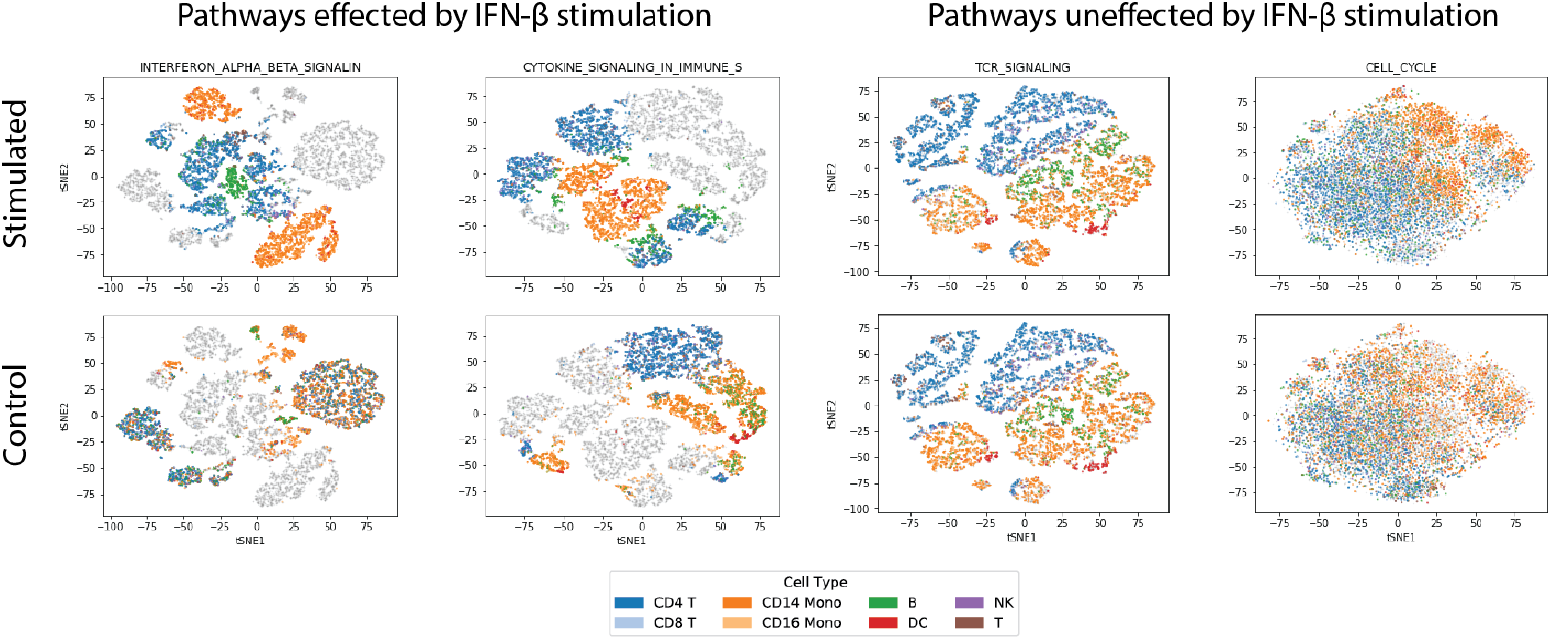
tSNE projections [50] computed from the 4-dimensional embeddings of the `Interferon-`*α*/*β*, `Cytokine signaling in immune system, TCR Signaling`, and `Cell cycle modules`. Each point represents a cell and is colored by its cell type, each column displays the projections of a single pathway, and cells from the control condition are greyed out in the top row, while cells from the stimulated condition are greyed out in the bottom row. We include two pathways targeted by the stimulation (`Interferon-`*α*/*β* and `Cytokine`) and two pathways not targeted (`TCR` and `Cell cycle`). These modules capture effects within the pathway contexts. For example, the perturbed `Interferon-`*α*/*β* and `Cytokine` modules strongly capture stimulation signals, while `TCR signaling`, which plays a role in cell-typing, captures cell-type specific effects. The `Cell cycle` is included as a control as we do not expect, nor observe, strong effects due to cell type or stimulus. More analysis of captured effects is discussed in Section 4.4.

We observe that only pathways directly related to the stimulation (`Interferon` *α/β* and `Cytokine signalling in the immune system`) are able to explain effects due to the stimulation, as evidenced by their ability to to discriminate between the stimulated (colored along the first row) and control (colored along the second row) cells. Furthermore, the module for the `TCR signaling pathway`, whose behavior is specific to cell-type context and is not involved in Interferon-*β* response, produces embeddings that capture cell-type specific contexts without capturing any stimulation effects. Finally, we observe that the control pathway, Cell cycle is unable to effectively discriminate either the stimulation status or the cell type, as we do not expect this pathway to be affected by either.

Interestingly, in the `Interferon` *α/β* representation, we find that the control cells without Interferon-*β* stimulus do not cluster by cell type, however, after stimulation we find strong clustering by cell type (Supplementary Figure 1). This behavior is not shared by its parent pathway `Cytokine signalling in the immune system`, whose control cells cluster by cell type, indicating that the response of `Interferon-`*α*/*β* is specific to its cell type context, a claim that is supported in the literature. van Boxel-Dezaire et al. [49] show that this cell type specific effect is caused by differences in the downstream transcription factors STAT1, STAT3, and STAT5 for CD4, CD8 T cells, B cells, and monocytes from human blood treated with Interferon-*β*. Additionally, this effect was also shown independently through a microarray experiment directly measuring differences between monocytes and T-cells after treatment with Interferon-*β* [15]. While modelling a context dependent response as observed in the Kang et al. dataset is difficult to accomplish in a univariate representation, it is easily captured in these multidimensional embeddings.

## 5 Discussion

In this paper, we presented pmVAE, a method to learn multidimensional pathway representations that are directly interpretable and reveal responses and effects that are specific to different pathway activities. By incorporating pathway membership into the architecture design, pmVAE constructs a latent space factorized by pathway. No further post-hoc analysis methods are required to identify such effects. This means that representations learned by pmVAE can be used to directly associate pathway states to clinically relevant features, quickly illuminating biological mechanisms underlying the data.

Through the use of our pathway module training procedure, we are able to address the problem of learning representations when pathways are highly overlapping, a complication innate to most pathway structures. We have empirically shown through the use of two scRNA-seq datasets containing biological noise from both direct stimulation, cell type, and in one example gRNA transduction, that our method produces biologically relevant and accurate pathway representations. We find that both pmVAE, and the one-dimensional variant of our model, 1D pmVAE, outperform baseline methods in being able to discriminate the cell state using the representation of the directly stimulated pathway. Additionally, we find that our learned representation of the true perturbed pathway is not only discriminative of the underlying biological signal, but that the closely related pathways, up- or downstream, are comparably discriminative.

We also demonstrate that our multidimensional pathway representations, as opposed to their unidimensional counterparts, are necessary to explain some pathway effects. For example, we observed that Interferon-*β* induces a cell-type dependent response in its target, `Interferon` *α*/*β*. While a unidimensional representation would struggle to explain such a context-dependent behavior, we find this effect clearly explained within the multidimensional pmVAE `Interferon` *α/β* `signaling` module. Though we have demonstrated that some pathways effects require multidimensional representations to be explained, it is likely that this is not true for all pathways. Proper model regularization, achieved through the 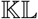 term and the tuning of its weight *β*, removes unnecessary complexity and effectively determines the correct dimensionality of each pathway module.

Redundancies between overlapping pathways, due in part to their hierarchical nature, result in degenerate solutions which make optimization challenging. To address this challenge, pmVAE enforces independence relationships between pathway modules by introducing local reconstruction terms for each module in the loss function. However, this independence ignores known pathway-pathway interactions arising from signaling effects. Learning pathway-factorized representations that explicitly model these effects, for example by incorporating known signaling interactions into the architecture [13, 22, 33, 40] is an interesting direction of future work. For example, after appropriate modeling of pathway-pathway interactions, remaining correlation could be attributed to sources of technical noise or bias, as can be done in mixed effect modelling.

Finally, although we have validated our method using only scRNA-seq data, we believe that our approach could work on other expression-based methods, such as bulk RNA-seq, CyTOF, DIA mass spectrometry, or any other technology that contain multiple measurements for each pathway of interest per sample. Through integration of multiple technologies one could be used to gain a more holistic understanding of cellular states. However, since these technologies might not have direct feature correspondences, the integrative analysis of them is challenging [18, 25]. While features sets might not be easily comparable, the pathway structure underlying these feature sets are shared. This indicates that pathway-factorized latent representations, like those learned by pmVAE, could be used more easily integrate [5,9,28,43] these technologies.

## References

[1] O. Acuto and F. Michel. CD28-mediated co-stimulation: a quantitative support for TCR signalling. Nat. Rev. Immunol., 3(12):939–951, Dec. 2003.

[2] M. Amodio, D. van Dijk, K. Srinivasan, W. S. Chen, H. Mohsen, K. R. Moon, A. Campbell, Y. Zhao, X. Wang, M. Venkataswamy, A. Desai, V. Ravi, P. Kumar, R. Montgomery, G. Wolf, and S. Krishnaswamy. Exploring single-cell data with deep multitasking neural networks. Nat. Methods, 16(11):1139–1145, Nov. 2019.

[3] J. Y. Bu, A. S. Shaw, and A. C. Chan. Analysis of the interaction of ZAP-70 and syk protein-tyrosine kinases with the t-cell antigen receptor by plasmon resonance. Proc. Natl. Acad. Sci. U. S. A., 92(11):5106–5110, May 1995.

[4] F. Buettner, N. Pratanwanich, D. J. McCarthy, J. C. Marioni, and O. Stegle. f-scLVM: scalable and versatile factor analysis for single-cell RNA-seq. Genome Biol., 18(1):212, Nov. 2017.

[5] K. Cao, X. Bai, Y. Hong, and L. Wan. Unsupervised topological alignment for single-cell multi-omics integration. Bioinformatics, 36(Supplement_1):i48–i56, 2020.

[6] D.-A. Clevert, T. Unterthiner, and S. Hochreiter. Fast and accurate deep network learning by exponential linear units (elus). arXiv preprint arXiv:1511.07289, 2015.

[7] P. Datlinger, A. F. Rendeiro, C. Schmidl, T. Krausgruber, P. Traxler, J. Klughammer, L. C. Schuster, A. Kuchler, D. Alpar, and C. Bock. Pooled crispr screening with single-cell transcriptome readout. Nature methods, 14(3):297–301, 2017.

[8] A. De La Hera, U. Müller, C. Olsson, S. Isaaz, and A. Tunnacliffe. Structure of the T cell antigen receptor (TCR): two CD3 epsilon subunits in a functional TCR/CD3 complex. J. Exp. Med., 173(1):7–17, 1991.

[9] P. Demetci, R. Santorella, B. Sandstede, W. S. Noble, and R. Singh. Gromov-wasserstein optimal transport to align single-cell multi-omics data. BioRxiv, 2020.

[10] A. B. Dincer, S. Celik, N. Hiranuma, and S.-I. Lee. Deepprofile: Deep learning of cancer molecular profiles for precision medicine. bioRxiv, page 278739, 2018.

[11] G. Eraslan, L. M. Simon, M. Mircea, N. S. Mueller, and F. J. Theis. Single-cell RNA-seq denoising using a deep count autoencoder. Nat. Commun., 10(1):390, Jan. 2019.

[12] A. Fabregat, S. Jupe, L. Matthews, K. Sidiropoulos, M. Gillespie, P. Garapati, R. Haw, B. Jassal, F. Korninger, B. May, et al. The reactome pathway knowledgebase. Nucleic acids research, 46(D1):D649–D655, 2018.

[13] N. Fortelny and C. Bock. Knowledge-primed neural networks enable biologically interpretable deep learning on single-cell sequencing data. Genome Biol., 21(1):190, Aug. 2020.

[14] K. He, X. Zhang, S. Ren, and J. Sun. Delving deep into rectifiers: Surpassing human-level performance on imagenet classification. In Proceedings of the IEEE international conference on computer vision, pages 1026–1034, 2015.

[15] N. Henig, N. Avidan, I. Mandel, E. Staun-Ram, E. Ginzburg, T. Paperna, R. Y. Pinter, and A. Miller. Interferon-beta induces distinct gene expression response patterns in human monocytes versus t cells. PloS one, 8(4):e62366, 2013.

[16] S. C. Hicks, F. W. Townes, M. Teng, and R. A. Irizarry. Missing data and technical variability in single-cell rna-sequencing experiments. Biostatistics, 19(4):562–578, 2018.

[17] J.-R. Hwang, Y. Byeon, D. Kim, and S.-G. Park. Recent insights of T cell receptor-mediated signaling pathways for T cell activation and development. Exp. Mol. Med., 52(5):750–761, May 2020.

[18] A. Irmisch, X. Bonilla, S. Chevrier, K.-V. Lehmann, F. Singer, N. Toussaint, C. Esposito, J. Mena, E. S. Milani, R. Casanova, et al. The tumor profiler study: integrated, multi-omic, functional tumor profiling for clinical decision support. medRxiv, 2020.

[19] H. M. Kang, M. Subramaniam, S. Targ, M. Nguyen, L. Maliskova, E. McCarthy, E. Wan, S. Wong, L. Byrnes, C. M. Lanata, et al. Multiplexed droplet single-cell rna-sequencing using natural genetic variation. Nature biotechnology, 36(1):89, 2018.

[20] D. P. Kingma and J. Ba. Adam: A method for stochastic optimization. arXiv, Dec. 2014.

[21] D. P. Kingma and M. Welling. Auto-Encoding variational bayes. arXiv, Dec. 2013.

[22] T. N. Kipf and M. Welling. Variational graph auto-encoders. arXiv preprint arXiv:1611.07308, 2016.

[23] B. Kompa and B. Coker. Learning a latent space of highly multidimensional cancer data. In Pacific Symposium on Biocomputing. Pacific Symposium on Biocomputing, volume 25, pages 379–390. World Scientific, 2020.

[24] B. M. Kuenzi, J. Park, S. H. Fong, K. S. Sanchez, J. Lee, J. F. Kreisberg, J. Ma, and T. Ideker. Predicting drug response and synergy using a deep learning model of human cancer cells. Cancer Cell, 38(5):672–684.e6, Nov. 2020.

[25] D. Lähnemann, J. Köster, E. Szczurek, D. J. McCarthy, S. C. Hicks, M. D. Robinson, C. A. Vallejos, K. R. Campbell, N. Beerenwinkel, A. Mahfouz, et al. Eleven grand challenges in single-cell data science. Genome biology, 21(1):1–35, 2020.

[26] C.-Y. Lee, S. Xie, P. Gallagher, Z. Zhang, and Z. Tu. Deeply-supervised nets. In Artificial intelligence and statistics, pages 562–570. PMLR, 2015.

[27] F. Letourneur and R. D. Klausner. Activation of T cells by a tyrosine kinase activation domain in the cytoplasmic tail of CD3 epsilon. Science, 255(5040):79–82, Jan. 1992.

[28] J. Liu, Y. Huang, R. Singh, J.-P. Vert, and W. S. Noble. Jointly embedding multiple single-cell omics measurements. BioRxiv, page 644310, 2019.

[29] F. Locatello, S. Bauer, M. Lucic, G. Raetsch, S. Gelly, B. Schölkopf, and O. Bachem. Challenging common assumptions in the unsupervised learning of disentangled representations. In international conference on machine learning, pages 4114–4124. PMLR, 2019.

[30] R. Lopez, J. Regier, M. B. Cole, M. I. Jordan, and N. Yosef. Deep generative modeling for single-cell transcriptomics. Nat. Methods, 15(12):1053–1058, Dec. 2018.

[31] M. Lotfollahi, F. A. Wolf, and F. J. Theis. scgen predicts single-cell perturbation responses. Nat. Methods, 16(8):715–721, Aug. 2019.

[32] M. D. Luecken and F. J. Theis. Current best practices in single-cell rna-seq analysis: a tutorial. Molecular systems biology, 15(6):e8746, 2019.

[33] J. Ma, M. K. Yu, S. Fong, K. Ono, E. Sage, B. Demchak, R. Sharan, and T. Ideker. Using deep learning to model the hierarchical structure and function of a cell. Nat. Methods, 15(4):290–298, Apr. 2018.

[34] W. Mao, E. Zaslavsky, B. M. Hartmann, S. C. Sealfon, and M. Chikina. Pathway-level information extractor (PLIER) for gene expression data. Nat. Methods, 16(7):607–610, July 2019.

[35] K. R. Moon, D. van Dijk, Z. Wang, S. Gigante, D. B. Burkhardt, W. S. Chen, K. Yim, A. van den Elzen, M. J. Hirn, R. R. Coifman, N. B. Ivanova, G. Wolf, and S. Krishnaswamy. Visualizing structure and transitions in high-dimensional biological data. Nat. Biotechnol., 37(12):1482–1492, Dec. 2019.

[36] S. Pestka, J. A. Langer, K. C. Zoon, and C. E. Samuel. Interferons and their actions. Annual review of biochemistry, 56(1):727–777, 1987.

[37] L. C. Platanias. Mechanisms of type-i- and type-II-interferon-mediated signalling. Nat. Rev. Immunol., 5(5):375–386, May 2005.

[38] S. Rybakov, M. Lotfollahi, F. J. Theis, and F. A. Wolf. Learning interpretable latent autoencoder representations with annotations of feature sets. bioRxiv, 2020.

[39] R. Satija, J. A. Farrell, D. Gennert, A. F. Schier, and A. Regev. Spatial reconstruction of single-cell gene expression data. Nat. Biotechnol., 33(5):495–502, May 2015.

[40] F. Scarselli, M. Gori, A. C. Tsoi, M. Hagenbuchner, and G. Monfardini. The graph neural network model. IEEE transactions on neural networks, 20(1):61–80, 2008.

[41] R. Schwanbeck, S. Martini, K. Bernoth, and U. Just. The notch signaling pathway: molecular basis of cell context dependency. European journal of cell biology, 90(6–7):572–581, 2011.

[42] N. Srivastava, G. Hinton, A. Krizhevsky, I. Sutskever, and R. Salakhutdinov. Dropout: a simple way to prevent neural networks from overfitting. The journal of machine learning research, 15(1):1929–1958, 2014.

[43] S. G. Stark, J. Ficek, F. Locatello, X. Bonilla, S. Chevrier, F. Singer, G. Rätsch, and K.-V. Lehmann. Scim: universal single-cell matching with unpaired feature sets. Bioinformatics, 36(Supplement_2):i919–i927, 2020.

[44] A. Subramanian, P. Tamayo, V. K. Mootha, S. Mukherjee, B. L. Ebert, M. A. Gillette, A. Paulovich, S. L. Pomeroy, T. R. Golub, E. S. Lander, and J. P. Mesirov. Gene set enrichment analysis: a knowledge-based approach for interpreting genome-wide expression profiles. Proc. Natl. Acad. Sci. U. S. A., 102(43):15545–15550, Oct. 2005.

[45] V. Svensson, A. Gayoso, N. Yosef, and L. Pachter. Interpretable factor models of single-cell rna-seq via variational autoencoders. Bioinformatics, 36(11):3418–3421, 2020.

[46] C. Szegedy, W. Liu, Y. Jia, P. Sermanet, S. Reed, D. Anguelov, D. Erhan, V. Vanhoucke, and A. Rabinovich. Going deeper with convolutions. In Proceedings of the IEEE conference on computer vision and pattern recognition, pages 1–9, 2015.

[47] J. Tan, M. Ung, C. Cheng, and C. S. Greene. Unsupervised feature construction and knowledge extraction from genome-wide assays of breast cancer with denoising autoencoders. In Pacific Symposium on Biocomputing Co-Chairs, pages 132–143. World Scientific, 2014.

[48] P.-Y. Tung, J. D. Blischak, C. J. Hsiao, D. A. Knowles, J. E. Burnett, J. K. Pritchard, and Y. Gilad. Batch effects and the effective design of single-cell gene expression studies. Scientific reports, 7:39921, 2017.

[49] A. H. van Boxel-Dezaire, J. A. Zula, Y. Xu, R. M. Ransohoff, J. W. Jacobberger, and G. R. Stark. Major differences in the responses of primary human leukocyte subsets to ifn-*β*. The Journal of Immunology, 185(10):5888–5899, 2010.

[50] L. Van der Maaten and G. Hinton. Visualizing data using t-sne. Journal of machine learning research, 9(11), 2008.

[51] D. van Dijk, R. Sharma, J. Nainys, K. Yim, P. Kathail, A. J. Carr, C. Burdziak, K. R. Moon, C. L. Chaffer, D. Pattabiraman, B. Bierie, L. Mazutis, G. Wolf, S. Krishnaswamy, and D. Pe’er. Recovering gene interactions from Single-Cell data using data diffusion. Cell, 174(3):716–729.e27, July 2018.

[52] G. P. Way and C. S. Greene. Extracting a biologically relevant latent space from cancer transcriptomes with variational autoencoders. Pac Symp Biocomput, 2018.

[53] G. P. Way, M. Zietz, V. Rubinetti, D. S. Himmelstein, and C. S. Greene. Compressing gene expression data using multiple latent space dimensionalities learns complementary biological representations. Genome Biology, 21(1):1–27, 2020.

[54] E. White. Deconvoluting the context-dependent role for autophagy in cancer. Nature reviews cancer, 12(6):401–410, 2012.

[55] F. A. Wolf, P. Angerer, and F. J. Theis. SCANPY: large-scale single-cell gene expression data analysis. Genome Biol., 19(1):15, Feb. 2018.

[56] C. Zhao, C. Denison, J. M. Huibregtse, S. Gygi, and R. M. Krug. Human isg15 conjugation targets both ifn-induced and constitutively expressed proteins functioning in diverse cellular pathways. Proceedings of the National Academy of Sciences, 102(29):10200–10205, 2005.

