## Supplemental Materials for "pmVAE: Learning Interpretable Single-Cell Representations with Pathway Modules"

January 28, 2021

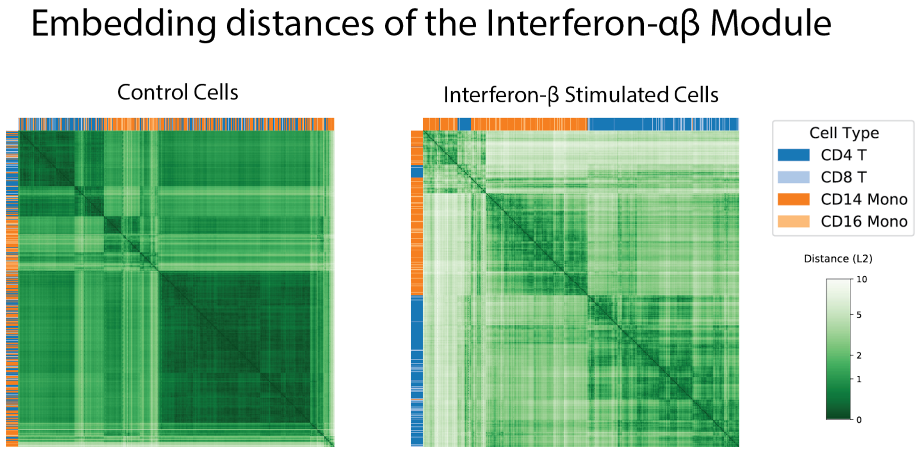


Figure 1: Heatmap analysis of T-Cells and Monocytes using the `Interferon-`$\alpha/\beta$ module. We observe that the control cells do not strongly cluster by cell type. However, after stimulation the cell types are mostly separated from one another. This suggests that the effect of the stimulation may be cell-type specific. This hypothesis is supported in the literature, where a similar effect has been shown for bulk RNA-sequencing [2].

| | median | max | min |
|---|---|---|---|
| **COSTIMULATION_BY_THE_CD28_FAMI** | 0.80 | 0.81 | 0.78 |
| **TCR_SIGNALING** | 0.78 | 0.80 | 0.77 |
| CELL_SURFACE_INTERACTIONS_AT_T | 0.76 | 0.79 | 0.72 |
| INTERFERON_GAMMA_SIGNALING | 0.75 | 0.77 | 0.73 |
| REGULATION_OF_MRNA_STABILITY_B | 0.74 | 0.76 | 0.73 |
| IMMUNOREGULATORY_INTERACTIONS_ | 0.74 | 0.76 | 0.72 |
| SEMAPHORIN_INTERACTIONS | 0.74 | 0.75 | 0.72 |
| INTERFERON_SIGNALING | 0.73 | 0.76 | 0.64 |
| APOPTOSIS | 0.73 | 0.74 | 0.72 |
| CELL_DEATH_SIGNALLING_VIA_NRAG | 0.73 | 0.75 | 0.72 |

Table 1: **pmVAE**: Top 10 pathways ranked by the median accuracy using logistic regression classifier for the Datlinger et. al. dataset [1], where the best and worst accuracy across 10 randomly seeded training runs are given through min and max. The bolded pathways are considered to be targeted by the anti-CD3 and the anti-CD28 antibodies. Median enrichment score using the top 10 pathways:0.014; test: hypergeometric.

| | median | max | min |
|---|---|---|---|
| APOPTOSIS | 0.76 | 0.78 | 0.74 |
| HEMOSTASIS | 0.73 | 0.78 | 0.53 |
| TRANSPORT_OF_INORGANIC_CATIONS | 0.72 | 0.73 | 0.71 |
| **TCR_SIGNALING** | 0.72 | 0.73 | 0.71 |
| IMMUNOREGULATORY_INTERACTIONS_ | 0.70 | 0.73 | 0.68 |
| CELL_DEATH_SIGNALLING_VIA_NRAG | 0.69 | 0.73 | 0.46 |
| INTERFERON_GAMMA_SIGNALING | 0.68 | 0.69 | 0.67 |
| TRANSMISSION_ACROSS_CHEMICAL_S | 0.68 | 0.69 | 0.65 |
| RESPONSE_TO_ELEVATED_PLATELET_ | 0.68 | 0.68 | 0.67 |
| NEURONAL_SYSTEM | 0.67 | 0.68 | 0.66 |

Table 2: **1D pmVAE**: Top 10 pathways ranked by the median accuracy using logistic regression classifier for the Datlinger et. al. dataset [1], where the best and worst accuracy across 10 randomly seeded training runs are given through min and max. The bolded pathways are considered to be targeted by the anti-CD3 and the anti-CD28 antibodies. Median enrichment score using the top 10 pathways:0.19; test: hypergeometric.

| | median | max | min |
|---|---|---|---|
| HEMOSTASIS | 0.78 | 0.86 | 0.55 |
| NEURONAL_SYSTEM | 0.74 | 0.80 | 0.70 |
| APOPTOSIS | 0.70 | 0.74 | 0.62 |
| **COSTIMULATION_BY_THE_CD28_FAMI** | 0.69 | 0.79 | 0.53 |
| TRANSMEMBRANE_TRANSPORT_OF_SMA | 0.69 | 0.81 | 0.55 |
| UNFOLDED_PROTEIN_RESPONSE | 0.67 | 0.71 | 0.57 |
| PHOSPHOLIPID_METABOLISM | 0.66 | 0.75 | 0.59 |
| DEVELOPMENTAL_BIOLOGY | 0.66 | 0.80 | 0.48 |
| **ADAPTIVE_IMMUNE_SYSTEM** | 0.66 | 0.86 | 0.56 |
| REGULATION_OF_MRNA_STABILITY_B | 0.65 | 0.78 | 0.49 |

Table 3: **Interpretable Autoencoder**: Top 10 pathways ranked by the median accuracy using logistic regression classifier for the Datlinger et. al. dataset [1], where the best and worst accuracy across 10 randomly seeded training runs are given through min and max. The bolded pathways are considered to be targeted by the anti-CD3 and the anti-CD28 antibodies. Median enrichment score using the top 10 pathways:0.19 ; test: hypergeometric.

| | median | max | min |
|---|---|---|---|
| GENERIC_TRANSCRIPTION_PATHWAY | 0.74 | 0.83 | 0.64 |
| MITOTIC_G1_G1_S_PHASES | 0.65 | 0.68 | 0.59 |
| L1CAM_INTERACTIONS | 0.64 | 0.71 | 0.61 |
| SEMAPHORIN_INTERACTIONS | 0.63 | 0.68 | 0.61 |
| NFKB_AND_MAP_KINASES_ACTIVATIO | 0.63 | 0.67 | 0.57 |
| HEMOSTASIS | 0.63 | 0.67 | 0.60 |
| CHROMOSOME_MAINTENANCE | 0.62 | 0.68 | 0.57 |
| UNFOLDED_PROTEIN_RESPONSE | 0.61 | 0.63 | 0.59 |
| S_PHASE | 0.59 | 0.69 | 0.54 |
| INNATE_IMMUNE_SYSTEM | 0.59 | 0.67 | 0.55 |

Table 4: **f-scLVM**: Top 10 pathways ranked by the median accuracy using logistic regression classifier for the Datlinger et. al. dataset [1], where the best and worst accuracy across 10 randomly seeded training runs are given through min and max. The bolded pathways are considered to be targeted by the anti-CD3 and the anti-CD28 antibodies. Enrichment score using the top 10 pathways: 0.8; test: hypergeometric.

| | median | max | min |
|---|---|---|---|
| **INTERFERON_SIGNALING** | 0.98 | 0.98 | 0.98 |
| IMMUNE_SYSTEM | 0.97 | 0.98 | 0.68 |
| **CYTOKINE_SIGNALING_IN_IMMUNE_S** | 0.97 | 0.98 | 0.97 |
| **INTERFERON_ALPHA_BETA_SIGNALIN** | 0.97 | 0.98 | 0.97 |
| INNATE_IMMUNE_SYSTEM | 0.97 | 0.97 | 0.96 |
| **ANTIVIRAL_MECHANISM_BY_IFN_STI** | 0.96 | 0.97 | 0.96 |
| RIG_I_MDA5_MEDIATED_INDUCTION_ | 0.96 | 0.96 | 0.95 |
| INTERFERON_GAMMA_SIGNALING | 0.90 | 0.90 | 0.87 |
| APOPTOSIS | 0.81 | 0.82 | 0.78 |
| TRIF_MEDIATED_TLR3_SIGNALING | 0.80 | 0.80 | 0.79 |

Table 5: **pmVAE**: Top 10 pathways ranked by the median accuracy using logistic regression classifier for the Kang et. al. dataset [3]. The max and the min columns are the best and worst accuracy across 10 randomly seeded training runs. The bolded pathways are considered to be targeted by interferon $\beta$ stimulation. Median enrichment score using the top 10 pathways: $1.6 \cdot 10^{-5}$; test: hypergeometric.

| | median | max | min |
|---|---|---|---|
| **INTERFERON_SIGNALING** | 0.97 | 0.98 | 0.72 |
| **INTERFERON_ALPHA_BETA_SIGNALIN** | 0.97 | 0.97 | 0.96 |
| **ANTIVIRAL_MECHANISM_BY_IFN_STI** | 0.96 | 0.97 | 0.96 |
| RIG_I_MDA5_MEDIATED_INDUCTION_ | 0.95 | 0.96 | 0.95 |
| APOPTOSIS | 0.82 | 0.82 | 0.81 |
| TRIF_MEDIATED_TLR3_SIGNALING | 0.78 | 0.80 | 0.78 |
| IMMUNE_SYSTEM | 0.66 | 0.97 | 0.44 |
| ACTIVATED_TLR4_SIGNALLING | 0.66 | 0.80 | 0.55 |
| SYNTHESIS_OF_DNA | 0.62 | 0.63 | 0.58 |
| METABOLISM_OF_CARBOHYDRATES | 0.62 | 0.63 | 0.61 |

Table 6: **1D pmVAE**: Top 10 pathways ranked by the median accuracy using logistic regression classifier for the Kang et. al. dataset [3]. The max and the min columns are the best and worst accuracy across 10 randomly seeded training runs. The bolded pathways are considered to be targeted by interferon $\beta$ stimulation. Median enrichment score using the top 10 pathways: $1.2 \cdot 10^{-3}$; test: hypergeometric.

| | median | max | min |
|---|---|---|---|
| **CYTOKINE_SIGNALING_IN_IMMUNE_S** | 0.96 | 0.98 | 0.91 |
| **INTERFERON_SIGNALING** | 0.92 | 0.95 | 0.50 |
| **INTERFERON_ALPHA_BETA_SIGNALIN** | 0.82 | 0.88 | 0.55 |
| RIG_I_MDA5_MEDIATED_INDUCTION_ | 0.82 | 0.91 | 0.74 |
| APOPTOSIS | 0.81 | 0.82 | 0.55 |
| **ANTIVIRAL_MECHANISM_BY_IFN_STI** | 0.74 | 0.80 | 0.44 |
| PURINE_METABOLISM | 0.66 | 0.67 | 0.57 |
| CELL_SURFACE_INTERACTIONS_AT_T | 0.64 | 0.68 | 0.53 |
| PEPTIDE_LIGAND_BINDING_RECEPTO | 0.63 | 0.80 | 0.51 |
| ACTIVATED_TLR4_SIGNALLING | 0.63 | 0.70 | 0.52 |

Table 7: **Interpretable Autoencoder**: Top 10 pathways ranked by the median accuracy using logistic regression classifier for the Kang et. al. dataset [3]. The max and the min columns are the best and worst accuracy across 10 randomly seeded training runs. The bolded pathways are considered to be targeted by interferon $\beta$ stimulation. Median enrichment score using the top 10 pathways: $1.2 \cdot 10^{-3}$; test: hypergeometric.

| | median | max | min |
|---|---|---|---|
| **CYTOKINE_SIGNALING_IN_IMMUNE_S** | 0.82 | 0.83 | 0.80 |
| GLYCOSAMINOGLYCAN_METABOLISM | 0.59 | 0.60 | 0.58 |
| BIOLOGICAL_OXIDATIONS | 0.57 | 0.59 | 0.56 |
| TRANS_GOLGI_NETWORK_VESICLE_BU | 0.56 | 0.59 | 0.54 |
| G_ALPHA_I_SIGNALLING_EVENTS | 0.56 | 0.57 | 0.56 |
| HEMOSTASIS | 0.55 | 0.56 | 0.54 |
| METABOLISM_OF_CARBOHYDRATES | 0.54 | 0.55 | 0.53 |
| SIGNALING_BY_GPCR | 0.54 | 0.56 | 0.53 |
| PURINE_METABOLISM | 0.54 | 0.55 | 0.52 |
| METABOLISM_OF_RNA | 0.54 | 0.56 | 0.53 |

Table 8: **f-scLVM**: Top 10 pathways ranked by the median accuracy using logistic regression classifier for the Kang et. al. dataset [3]. The max and the min columns are the best and worst accuracy across 10 randomly seeded training runs. The bolded pathways are considered to be targeted by interferon $\beta$ stimulation. Enrichment score using the top 10 pathways: 0.24; test: hypergeometric.

### References

[1] P. Datlinger, A. F. Rendeiro, C. Schmidl, T. Krausgruber, P. Traxler, J. Klughammer, L. C. Schuster, A. Kuchler, D. Alpar, and C. Bock. Pooled crispr screening with single-cell transcriptome readout. *Nature methods*, 14(3):297–301, 2017.

[2] N. Henig, N. Avidan, I. Mandel, E. Staun-Ram, E. Ginzburg, T. Paperna, R. Y. Pinter, and A. Miller. Interferon-beta induces distinct gene expression response patterns in human monocytes versus t cells. *PloS one*, 8(4):e62366, 2013.

[3] H. M. Kang, M. Subramaniam, S. Targ, M. Nguyen, L. Maliskova, E. McCarthy, E. Wan, S. Wong, L. Byrnes, C. M. Lanata, et al. Multiplexed droplet single-cell rna-sequencing using natural genetic variation. *Nature biotechnology*, 36(1):89, 2018.
